# Macronutrient-preference is modulated by biological sex and estrous cycle in mice

**DOI:** 10.64898/2026.03.26.714595

**Authors:** A. Dofat, R. Jacob, K. Jacobs, M. Ahrens, W.M. Howe

## Abstract

Dietary choice plays a critical role in metabolic and neurological health, yet the biological factors that shape macronutrient preference remain poorly understood. Evidence from both humans and rodents suggests potential sex differences in the attractiveness of specific nutrients, though findings have been inconsistent and often rely on self-report or diets with mixed macronutrient composition. The present study examined sex differences in macronutrient preference and food-directed behavior in mice using a controlled three-food choice paradigm. Adult male (n = 12) and female (n = 11) C57BL/6J mice were given simultaneous access to foods consisting of fat, sucrose, or a fat–carbohydrate combination across 14 days. Intake, latency to approach, and time spent near each food source were quantified, and estrous cycle stage was monitored in females. Female mice consumed significantly more food than males overall, driven by a selective increase in fat intake. Behavioral measures paralleled these results, with females spending more time in proximity to fat-associated food zones. In contrast, males preferentially consumed the fat–carbohydrate combination and showed weaker nutrient-specific engagement. Estrous cycle stage modestly influenced feeding behavior, with estrus associated with increased overall intake and greater consumption of combination diets, reflecting elevated carbohydrate intake. These findings demonstrate robust sex differences in macronutrient preference and suggest that hormonal state may selectively modulate nutrient-specific feeding behavior.

## Introduction

Diet modifies the risk of cardiovascular, metabolic, and neurodegenerative diseases and represents a major modifiable determinant of disease ^1–5^. Many factors can contribute to the decision-making process behind food selection. At the level of the individual, homeostatic drives can lead individuals to seek out different food sources based upon physiological needs. In the modern environment, enriched with easily accessible sources of calories, food choice is often guided by the subjective rewarding qualities of different foods, or preference. Prior studies have highlighted marked individual variability in food preferences, including potential biological sex differences in the attractiveness of different macronutrients. For example, some studies report that relative to men, women consume a higher proportion of energy from dietary fat even after controlling for differences in body weight ^6^. Indeed, even though women self-report more diet-conscious approaches and consume more fruits and vegetables compared to men ^7,8^, rates of obesity are higher in females perhaps due in part to greater fat consumption ^9–13^. In contrast, studies in mice have suggested that males exhibit a preference for carbohydrates ^14,15^. Recent research has begun to map out the neural circuit mechanisms that mediate the rewarding qualities of different macronutrients, highlighting key differences in the peripheral afferent systems that relay information about the macronutrient content of foods to the brain ^16–18^. Whereas fat reward depends upon PPAR-alpha mediated recruitment of vagal afferents to the hindbrain, carbohydrate (glucose) reinforcement signals depend upon the recruitment of hepatic portal vein sensors and spinal afferent pathways ^16,18,19^. Critically, research has also highlighted sex differences in peripheral fat and glucose sensing and metabolic systems ^20–24^, suggesting that the above descriptions of sex differences in macronutrient preference may stem from fundamental differences in the organization of peripheral-central metabolic pathways. Additionally, sex hormones, particularly estrogen, have been shown to influence food cravings and overall food intake ^25–29^, and fluctuations in estrogen levels drive cyclical changes in taste preferences and cravings, particularly for sweet and energy-dense foods ^30–34^. However, it is important to note that findings regarding differences in food preference between males and females are inconsistent ^35^. Much of the existing literature from humans relies heavily on self-reported data, which can introduce bias and limit the accuracy of findings. Food categories in these studies are often inconsistently defined, making it difficult to draw precise conclusions about macronutrient-specific differences. Many human studies also focus on food cravings ^30–35^, though cravings do not always directly translate to food intake. Similarly, work in model systems like rodents often juxtapose intake of standard lab chow or sucrose pellets against commercially available “high-fat” diets, which are typically combinations of fats are carbohydrates that may have unique effects on brain reward systems that magnify their attractiveness, and thereby confound conclusions about fat or sugar preferences *per se* ^17,36^. These limitations highlight the need for more controlled experiments designed to probe for underlying sex differences in macronutrient-specific food preference. To address these gaps, here we assessed macronutrient preference in male and female mice. Given the potential for modulation of preference by circulating hormone levels, we systematically tracked intake and estrous cycle across 14 days.

## Methods

### Animals

Twenty-three Adult (3 - 4-month-old) male and female C57BL/6J (n = 12 male and n = 11 female) mice were acquired from Jackson Laboratories (Bar Harbor, ME, USA) were housed in colony cages with free access to standard lab chow (Teklad Global 18% Protein Rodent Diet; Envigo, Indianapolis, IN, USA; product no. 2018) and water. Animals were maintained on a 12-hour reverse light/dark cycle, with behavioral experiments conducted in the dark phase. All procedures used in the present study were reviewed and approved by the Animal Care and Use Committee at Virginia Tech and conformed to National Institutes of Health (NIH) guidelines. On the first day of the experiment, animals were weighed and placed on a mild food deprivation schedule. Food was removed from their cages to reduce their body weight to 90–95% of their free-feeding weight, which was maintained for the entirety of the experiment.

### Tracking Estrous cycle

Vaginal lavage was performed after each habituation and experimental recording day to determine the estrous cycle stage of the female mice. A sterile 200 μL pipette tip attached to a pipette was used to draw approximately 100 μL of sterile saline solution. The mouse was placed on the cage topper, and the tail was gently lifted to elevate the rear end. The end of the pipette tip was placed at the opening of the vaginal canal. Approximately 25–50 μL of the saline solution was gently expelled at the opening, allowing the liquid to spontaneously flow into the canal. The pressure on the pipette was slowly released to withdraw the fluid back into the tip, avoiding rapid release to prevent aspiration of fluid into the pipette. This process was repeated 4–5 times using the same tip and fluid to collect enough cells in a single sample ^37^. The collected fluid was placed onto glass slides and allowed to air dry completely at room temperature. Once dry, the slides were immediately washed with 95% and 70% ETOH, 0.01M Sodium bicarbonate buffer, and stained using 1% Toluidine Blue. Estrous cycle stage was determined via vaginal cytology: proestrus was identified by the predominance of nucleated epithelial cells, estrus by the predominance of cornified epithelial cells, and diestrus by the predominance of leukocytes ^38^. For analysis, proestrus and estrus phases were combined and categorized as “estrus” based on both stages representing periods of elevated estrogen and maximal sexual receptivity in female mice.

### Food types

Three food types made of either fat, carbohydrate, or a blend of fat and carbohydrate were used in this study. The fat stimulus was prepared using Crisco® vegetable shortening (The J.M. Smucker Company, Orrville, OH, USA). The carbohydrate stimulus was chocolate-flavored sucrose pellets (LabTab™ AIN-76A Rodent Chocolate Tablets, 20 mg; TestDiet, Richmond, IN, USA). The last food type was a combination of the fat and carbohydrate (‘combo’) stimuli, created by grinding the carbohydrate pellets and mixing them with Crisco® vegetable shortening to achieve 1:1 caloric ratio between carbohydrate to fat; in other words, mice received the same number of calories from fat and carbohydrate per gram of ‘combo’ consumed.

### Three Food Choice preference test

Food preference was assessed using a Three Food Choice (3FC) paradigm (Figure 1A). The 3FC experiment took place in a 12 × 12 plexiglass chamber equipped with video tracking (Noldus Phenotyper). Training began following the initiation of food deprivation and after a 6-day habituation period to both the testing arena and the food. For habituation, each mouse was placed in the arena for 45 minutes daily and allowed to freely explore. At the completion of such chamber habituations, mice were provided with 1kCal of each of the above food types in their home cages. Each food type was provided alone for two consecutive days. On Day 7, three food dishes were placed in three of the four corners of the arena. Each dish contained 1 gram of either fat, carbohydrate, or combo. The food was weighed before and after each recording session to track food consumption. The locations of the foods were varied between mice to counterbalance any positional bias within the chambers and remained constant across days. At testing onset, mice were individually placed in the center of a behavioral arena, and their activity was video recorded for 45 minutes before being returned to their home cages. Food intake was measured over a consecutive 14-day testing period, after which animals were returned to ad libitum access to standard chow. Motion tracking video recordings were analyzed to additionally quantify the duration spent in proximity to each food zone and the latency to first approach each zone. Food zones were defined as regions surrounding each food bowl. A mouse was considered to be within a food zone when its head remained in the designated area for at least 2 seconds. Total time spent in each zone was calculated as the cumulative duration the mouse’s head was detected in each of the three food zones. Latency to approach each food type was defined as the time (in seconds) from the start of the recording corresponding to the mouse’s initial placement in the center of the arena until its first entry into each food zone.

**Figure 1.**
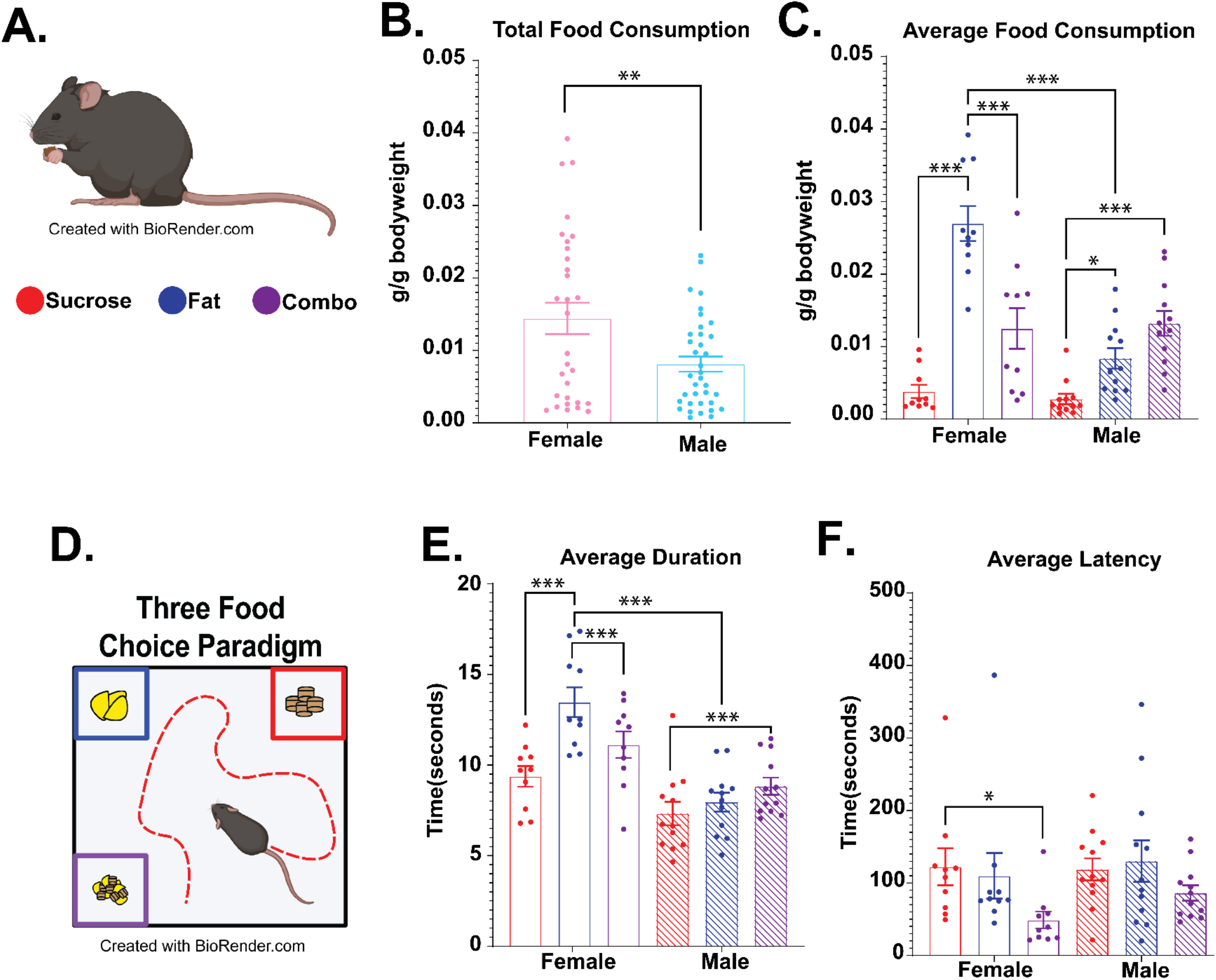
Biological sex differences in macronutrient intake and food-directed behavior. (A) Example image of mouse eating sucrose pellet. (B) Total food consumption normalized to body weight (g/g) across all food types for female and male mice. (C) Average food consumption by food type (sucrose, fat, combination) separated by sex. (D) Schematic of mouse exploring during the three-food choice assay in which mice were given simultaneous access to sucrose, fat, and a combination food source within a behavioral arena. (E) Average duration spent in proximity to each food zone. Overall, mice spent more time near fat and combination foods than sucrose. (F) Average latency to first approach each food zone. Data are shown as mean ± SEM with individual data points representing single animals. *p < 0.05, **p < 0.01, ***p < 0.001. Colors denote food type: sucrose (red), fat (blue), and combination (purple). Solid bars represent females and hatched bars

### Statistical Analysis

Behavioral data were acquired using EthoVision software and are presented as mean ± standard error of the mean (SEM). All analyses were conducted in R (Core Team, 2023 version 4.4.2). Food consumption (g food/g bodyweight and kcal/g), latency to approach, and duration near food bowls were averaged across week 1 and week 2 (Week 1: Mean Days 1–7; Week 2: Mean Days 8–14) within each mouse for each combination of food type (sucrose, fat, combination) and test week. Data were analyzed using linear mixed-effects models (LME) fit with the lme4 package (Bates et al., 2015) with intake, total time spent around each food (duration) and time to first approach each food (latency) as the outcome and sex, food type, and their interaction as fixed effects and mouse as a random intercept to account for repeated measures. Type III ANOVAs with Kenward–Roger approximation of degrees of freedom were used to test for overall significance for each covariate. Estimated marginal means (EMMs) and post hoc comparisons were conducted using the emmeans package (Lenth, 2023). Post hoc comparisons were performed on EMMs using Tukey adjustment to control family-wise error rate. Separate analyses were conducted to examine the influence of estrous cycle stage on behavior. Female data were segmented based on vaginal cytology into diestrus and averaged within each mouse Additional paired t-tests compared overall intake between diestrus and estrus. All statistical tests were two-tailed, and α was set at α = 0.05.

## Results

### Biological Sex Effect on Food Intake

When intake was normalized to body weight (g/g), we noted that food intake varied as a function of food type (F(2, 100) = 71.72, p < 0.01). Post hoc comparisons showed that, overall, fat intake exceeded both combination (t(108) = −2.40, p = 0.04) and sucrose (t(108) = 8.69, p < 0.01), and combination exceeded sucrose (t(108) = 6.29, p < 0.01). We also noted that overall food types of females consumed nearly double the amount of food compared to males (**Fig 1b**, main effect of biological sex (F(1, 20) = 20.49, p < 0.01 females 0.61 ± 0.61 g/g, males 0.30 ± 0.34g/g). Furthermore, males and females differed in the amount of each food type they consumed (Food Type × Sex interaction, F(2, 100) = 36.19, p < 0.01). Female mice showed a strong preference for fat, consuming more fat than both combination (t(48) = 6.16, p < 0.01) and sucrose (t(48) = 10.21, p < 0.01), and more combination than sucrose (t(48) = 4.05, p < 0.01; **Fig. 1c**). Similar to females, male mice consumed the least amount of sucrose (fat vs sucrose, t(58) = 4.81, p < 0.01; combo vs sucrose, t(58) = 8.83, p < 0.01). In contrast to females, males, consumed more combination than fat (**Fig 1c**, t(58) = 4.02, p < 0.01). Direct comparisons between sexes confirmed that females consumed significantly more fat than males (t(66.5) = 9.23, p < 0.01), while no sex differences were observed for combination (t(66.5) = −0.10, p =1.00) or sucrose intake (t(66.5) = 0.52, p = 0.99). Intake remained stable across time, with no main effect of Week (F(1, 100) = 1.19, p =0.22) and no significant interactions involving Week (all p > 0.27). These results suggest that the elevated food intake observed in females is primarily driven by a selective and stable elevation in dietary fat consumption.

### Macronutrient and sex effects on food interaction

The above analyses focused upon overall intake of each of the different food types. We next assessed the possibility that macronutrient composition or biological sex may influence the duration of time spent in proximity to each food type, as well as the latency for mice to first approach each food. The results of our analysis on duration largely echoed the general patterns of intake described above. We detected statistically significant main effects of Biological Sex (F(1, 20) = 28.14, p < 0.01), Week (F(1, 100) = 6.10, p =0.01), and Food type (F(2, 100) = 16.62, p < 0.01) on the total time spent around the different food stimuli. Post hoc comparisons indicated that mice spent significantly more time near fat (t(108) = 4.75, p < 0.01) and combination (t(108) = 3.47, p < 0.01) food zones than sucrose, with no difference between fat and combination (t(108) = 1.27, p =0.41 **Fig. 1d**). Females spent more time in food zones overall than males, averaging 475.55 ± 11.32 seconds compared to 337.71 ± 8.04 seconds in males (**Fig. 1i**). We noted that males and females also differed in the amount of time they spent around each food type (Food type × Sex (F(2, 100) = 10.60, p < 0.01). To examine sex-specific patterns of engagement, duration was analyzed separately by sex. In males, time spent near food zones varied by Food type (F(2, 58) = 5.48, p <0.01). Post hoc comparisons revealed greater duration near combination compared to sucrose (t(58) = 3.30, p < 0.01), but no differences between combination and fat (t(58) = 1.91, p = 0.14) or fat and sucrose (t(58) = 1.39, p =0.35; **Fig. 1d**). Females showed a distinct pattern (F(2, 48) = 14.10, p < 0.01), spending significantly more time near fat than combination (t(48) = 3.03, p =0.01) or sucrose (t(48) = 5.29, p < 0.01). Duration near combination was also numerically greater than sucrose but did not reach significance (t(48) = 2.26, p = 0.07), suggesting a selective enhancement of engagement by fat-rich foods. Direct sex comparisons confirmed that females spent more time near fat than males (t(50.5) = 6.93, p < 0.01), with no significant differences for combination (t(50.5) = 2.88, p = 0.06) or sucrose (t(50.5) = 2.57, p =0.12; **Fig. 1d**). Additionally, time spent in proximity to each food also varied across weeks of recording (F(1,100) = 6.10, p =0.01), however, this change in time spent around was sex-specific (Sex × Week (F(1, 100) = 5.81, p =0.01). Follow-up analyses indicated that time spent in proximity to the different foods was consistent across weeks in males (F(1, 59) = 0.0027, p =0.96), whereas females spent more time engaged with foods during Week 2 than Week 1 (F(1, 49) = 4.96, p =0.03). All other interactions were not significant (all p’s > 0.64).

With respect to latency to approach each food, we noted significant main effects of Food Type (F(2, 100) = 6.90, p < 0.01) and Week (F(1, 100) = 5.72, p =0.01). Post hoc comparisons indicated that mice approached the combination food zone significantly faster than either fat (t(108) = −3.10, p < 0.01) or sucrose (t(108) = −3.07, p < 0.01), while latencies for fat and sucrose did not differ (t(108) = 0.03, p = 0.99; **Fig. 1e**). We also noted that approach latency decreased over time, with all mice entering food zones more quickly in Week 2 than in Week 1. On average, latency was reduced by 30.67 seconds from Week 1 to Week 2 (t(21) = 2.50, p = 0.021). No main effect of Biological Sex was observed (F (1, 20) = 0.80, p = 0.38), and no interactions reached significance (all p > 0.23), indicating that sex did not influence approach latency. Combined, our results show that food-directed behavior was shaped by food type, sex, and experience. While males engaged most with combination foods. Even though they approached combination foods more readily than fat or sucrose only, females preferentially consumed fat. This increase in fat consumption was mirrored by an escalating amount of time spent around fat across training days.

### Estrous Cycle Effect on Food Intake

To evaluate the effects of estrous cycle stage on food intake and food-directed behavior, female data were reorganized such that test week was no longer included as a grouping factor. For each food type, intake and behavioral measures were averaged separately across all test days in which each mouse was classified as being in estrus or diestrus. Data were analyzed using a linear mixed-effects model with Food Type (sucrose, fat, combination) and Cycle Stage as fixed effects, and Mouse ID included as a random intercept. When intake was normalized to body weight (g/g), results paralleled those described above for female data, with a robust main effect of Food Type on intake (F(2, 324.09) = 125.11, p < 0.01). Furthermore, females consumed significantly more total food during estrus compared with diestrus (F(1, 211.12) = 4.17, p = 0.04; **Fig. 2d**). In addition, results showed that estrous cycle effects on intake were food type–specific (Cycle Stage × Food Type interaction, F(2, 324.09) = 3.89, p = 0.02). Post hoc comparisons revealed that females consumed significantly more combination food during estrus than during diestrus (t(323) = −3.17, p = 0.02), whereas fat and sucrose intake did not differ significantly between cycle stages (all p > 0.81; **Fig. 2e**). These findings suggest that the influence of the estrous cycle on food intake is driven primarily by increased consumption of combination food during estrus.

**Figure 2.**
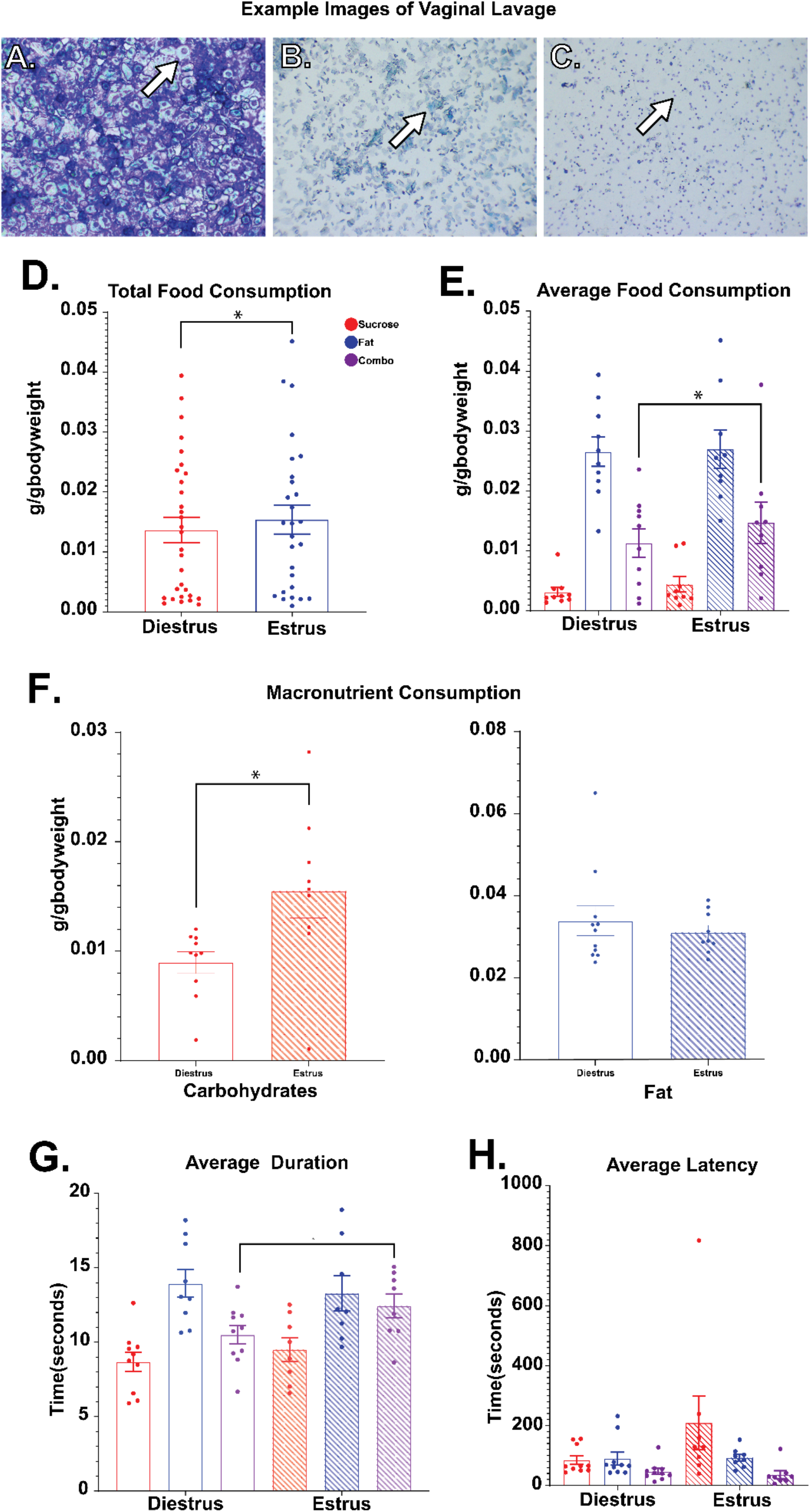
Estrous cycle modulation of macronutrient intake and food-directed behavior. (A–C) Representative vaginal lavage cytology images used to classify estrous cycle stage and each arrow indicated the cell type that identifies each stage (A: proestrus, nucleated epithelial cell, B: estrus, cornified epithelial cell, C: diestrus, leukocyte). (D) Total food consumption normalized to body weight (g/g) across all food types during diestrus and estrus. (E) Average food consumption by food type (sucrose, fat, combination) across estrous cycle stages. (F) Estimated macronutrient consumption (carbohydrates and fat) normalized to body weight (g/g) during diestrus and estrus. Macronutrient intake was calculated by accounting for the nutrient composition of each diet. (G) Average duration spent near each food type during diestrus and estrus. (H) Average latency to first approach each food zone across estrous cycle stages. Data are shown as mean ± SEM with individual data points representing single animals. Colors denote food type: sucrose (red), fat (blue), and combination (purple). Solid bars represent diestrus and hatched bars represent estrus. *p < 0.05, **p < 0.01, ***p < 0.001.

To further assess the influence of the estrous cycle on food preference, we examined whether total macronutrient intake differed between cycle stages. Total daily carbohydrate and fat intake (g/g body weight) was estimated by accounting for the nutrient composition of each diet. Because the combination diet contained 72.97% sucrose and 27.03% fat by weight, intake from this diet was partitioned accordingly and added to the respective totals from sucrose pellets and fat. This approach provided an estimate of the total amount of each macronutrient consumed during diestrus and estrus. Data was analyzed using a linear mixed-effects model with Macronutrient Type (carbohydrate, fat) and Cycle Stage as fixed effects and Mouse ID as a random intercept. Similar to the above cycle g/g data, total consumption differed between stages (F(1, 192.52) = 5.96, p = 0.01) and between macronutrient types (F(1, 212.35) = 120.17, p < 0.01). Additionally, the amount of each macronutrient type consumed was influenced by estrous cycle stage (Cycle Stage × Macronutrient Type interaction F(1, 212.35) = 5.28, p = 0.02) which was driven by increased carbohydrate intake during estrus seen in Post hoc comparisons that showed carbohydrate intake was significantly higher during estrus compared with diestrus (t(219) = −3.31, p < 0.01. However fat intake did not differ between stages (t(219) = −0.25, p =0.99; **Fig. 2f**). These findings indicate that estrous cycle–related changes in nutrient consumption are attributable to selective increases in carbohydrate intake during estrus.

### Food and estrous cycle effects on food interaction

While the above analyses focused on intake of each food and macronutrient type across the estrous cycle, we next examined whether cycle stage influenced food-directed behaviors, including time spent near each food type and latency to approach. Using the same linear mixed-effects model described above, we found that approach latency differed significantly by Food Type (F(2, 324.09) = 8.28, p < 0.01), but overall latency did not differ between the stages (F(1, 213.48) = 1.38, p = 0.23). Additionally, within each food type, latency didn’t differ between estrous or diestrus (Cycle Stage × Food Type interaction F(2, 324.09) = 2.14, p = 0.11; **Fig. 2i**).

In contrast, analysis of time spent near food zones revealed that the time spent around all food zones differed between estrus and diestrus (F(1, 323.17) = 5.86, p = 0.01; **Fig. 2g**) and between each food type (F(1, 324.01) = 40.77, p < 0.01). Additionally, there was also a difference in time spent around each food type due to cycle stage (Cycle Stage × Food Type interaction, F(1, 324.01) = 3.21, p = 0.04), indicating that the effect of estrous cycle on food engagement varied by food type. This effect was primarily driven by the increase in time females spent near combination food zones during estrus compared with diestrus (t(333) = −3.48, p < 0.01), whereas time spent near fat (t(333) = −0.34, p = 0.99) or sucrose (t(333) = −0.76, p = 0.97) zones did not differ between stages (**Fig. 2h**). These findings suggest that estrous cycle–related changes in food-directed behavior are specific to combination foods, which may be particularly salient or reinforcing during high-estrogen states.

## Discussion

The present study employed a three-choice feeding paradigm to investigate how biological sex and estrous cycle stage influence food type preference and food-directed behavior in mice across a 14-day testing period. Our findings revealed consistent and robust sex differences in food intake: female mice consumed significantly more food by weight (g/g) than males, an effect driven specifically by increased consumption of dietary fat. This pattern mirrors findings from human studies in which women report greater fat preference and obtain a higher proportion of calories from fat relative to men, even after controlling for body weight ^6^. This sex difference was food type-specific. Post hoc comparisons confirmed that female mice consumed more fat than males (p < 0.0001), but no sex differences were observed for sucrose or combination diets. Within females, fat intake significantly exceeded both sucrose and combination foods, while males showed a distinct pattern, consuming more combination than fat or sucrose. These data suggest that elevated intake in females reflects a selective preference for fat, instead of an overall increase in eating.

Behavioral measures further supported a sex-specific fat preference. Females spent more time in fat-associated food zones relative to other diets and relative to males, despite showing no differences in initial approach latency. This suggests that fat did not elicit stronger anticipatory behavior, but rather promoted sustained engagement after exposure, potentially reflecting a post-ingestive motivational mechanism. In contrast, males spent more time near combination food zones than sucrose but did not significantly differentiate between fat and sucrose. Latency and duration measures in males did not vary by food type, suggesting a lack of nutrient-specific motivation or reduced sensitivity to dietary cues in this context. Time-dependent changes also diverged by sex. While females maintained stable intake and behavioral engagement across the two-week period, males exhibited a significant increase in fat intake from Week 1 to Week 2. This effect was specific to fat and was not observed for sucrose or combination diets. These findings align with prior research indicating that nutrient-specific learning may be shaped by post-ingestive feedback (e.g., via vagal or dopaminergic signaling) and suggest that males may acquire a preference for fat over time, whereas females display an early and stable fat bias.

When examining nutrient-specific intake, estrous cycle stage exerted a modest but significant influence on total food consumption, with females consuming more food during estrus than diestrus. Importantly, this effect was driven specifically by increased intake of the combination diet, whereas consumption of fat-only or sucrose-only diets did not differ between cycle stages. Consistent with this pattern, analysis of total macronutrient intake revealed that estrus was associated with greater carbohydrate consumption, while fat intake remained unchanged. These findings suggest that estrous-related increases in intake may emerge preferentially in the context of foods that contain carbohydrates.

Estrous cycle stage also influenced food-directed behavior. Females spent more time near food zones during estrus compared with diestrus, and this effect was similarly driven by increased engagement with the combination food. In contrast, approach latency did not differ between cycle stages, suggesting that estrous effects were not attributable to generalized changes in exploratory behavior or motivation to initiate feeding. Instead, these results indicate that estrous cycle stage selectively enhances engagement with nutritionally complex foods once they are encountered. Together, these findings suggest that ovarian hormone fluctuations may modestly modulate nutrient selection and food-directed behavior, particularly in the context of foods containing mixed macronutrient compositions.

Collectively, these findings support emerging evidence that nutrient-specific gut-brain signaling pathways, particularly those related to fat sensing, such as PPAR-α activation and vagal afferents. may be differentially tuned by sex. Female-biased increases in fat consumption and food zone engagement may reflect enhanced sensitivity to fat-mediated post-ingestive signals, whereas the delayed fat preference observed in males may involve reinforcement learning mechanisms. These findings have important implications for understanding sex differences in vulnerability to diet-induced obesity and metabolic disorders, and underscore the need for sex-informed frameworks in nutritional neuroscience.

## Limitations

While this study provides novel insights into the sex- and hormone-related modulation of food type preference, several limitations should be noted. First, estrous cycle stage was assessed using vaginal cytology, a widely used but indirect proxy for hormonal status. Transitional phases or ambiguous cytological profiles may obscure subtle hormone effects. More precise control of hormonal state could be achieved through ovariectomy and hormone replacement approaches, enabling direct tests of estrogen and progesterone contributions to nutrient-specific intake. Second, this study employed voluntary oral feeding paradigms, which preserve ecological validity but introduce potential confounds related to orosensory cues. Palatability and taste sensitivity may differ by sex or estrous stage, influencing intake independently of post-ingestive mechanisms. Intragastric infusion studies could help disentangle these effects and directly assess the role of gut nutrient sensing. Third, the findings were derived from healthy adult mice under ad libitum feeding conditions. It remains unclear whether similar patterns would emerge under conditions of metabolic stress, caloric restriction, or obesity. Future work should explore how developmental stage, metabolic state, and dietary history interact with sex and hormonal state to shape nutrient preferences. Finally, while latency and duration metrics offer valuable behavioral endpoints, they are indirect proxies for motivational processes. Incorporating neural circuit mapping or optogenetic manipulations, or targeted assays of gut-brain signaling would help elucidate the underlying mechanisms through which sex and hormone state modulate food-directed behavior.

## Conclusion

This study demonstrates that biological sex exerts a significant influence on food type preference and food-directed behavior in mice. Female mice showed a consistent and selective preference for dietary fat, reflected in both intake and engagement metrics, while male mice exhibited a delayed shift toward fat preference over time. Estrous cycle stage had modest effects, primarily influencing intake of combination diets, but did not significantly alter overall latency. These results highlight sex-specific differences in nutrient valuation and motivation, likely mediated by divergent gut-brain signaling pathways, and underscore the importance of incorporating sex and hormonal state as key biological variables in studies of feeding behavior and metabolic health.

